# Patch-clamp recordings in slices of telencephalon, diencephalon and rhombencephalon of salamanders

**DOI:** 10.1101/2020.06.10.143487

**Authors:** Aurélie Flaive, Dimitri Ryczko

## Abstract

The salamander is a key limbed vertebrate from which many major scientific questions can be addressed in the fields of motor control, evolutionary biology, and regeneration biology. An important gap of knowledge is the description of the electrophysiological properties of the neurons constituting their central nervous system. To our knowledge, some patch-clamp electrophysiological recordings were done in the spinal cord and recently in hindbrain slices, but not in any higher brain region. Here, we present a method to obtain patch-clamp recordings in slices of the telencephalon, diencephalon and rhombencephalon of salamanders. The method includes dissection of the brain, brain slice preparation, visual identification of neurons and patch-clamp recordings. We provide single cell recordings in the rhombencephalon, diencephalon and telencephalon of salamanders. This method should open new avenues to dissect the operation of salamander brain circuits at the cellular level.

**Highlights:**

- Salamander brain slices of telencephalon, diencephalon, and rhombencephalon
- Patch-clamp recordings in salamander brain slices
- The salamander as a model to decipher tetrapod neural microcircuits

## 1. Introduction

Among limbed vertebrates, the salamander is a unique animal model to decipher the organisation of the locomotor neural circuitry, but also its evolution and its regeneration after major lesions. Salamanders swim underwater and walk on land, therefore providing an opportunity to dissect the interactions between the neural circuits controlling axial movements and those controlling limb movements (Ryczko et al. 2015). They are the closest representative of the first tetrapods, therefore allowing researchers to infer the evolution of the locomotor circuitry during the transition to land (Ijspeert et al. 2007). They regenerate their spinal cord after a complete transection (Chevallier et al. 2004) or after major destruction of their brain dopaminergic system (Joven et al. 2018), providing a unique opportunity to dissect the reconnection maps leading to functional recovery in limbed vertebrates.

The available knowledge of the electrophysiological properties of central salamander neurons is limited compared to other models. Mainly extracellular electrophysiological recordings were obtained from axial ventral roots or limb nerves during fictive locomotion (Ryczko et al. 2015), or from reticular nuclei following stimulation of the Mesencephalic Locomotor Region, a brainstem region that controls locomotion in vertebrates (Ryczko et al. 2016a). Sharp intracellular recordings were done during fictive locomotion in spinal circuits controlling limb (Wheatley and Stein 1992) or axial movements (Perrins and Soffe 1996). Patch-clamp recordings were successfully used to describe the cholinergic modulation of limb motoneuron activity in spinal cord slices (Chevallier et al. 2006). At the brain level, only one study used patch-clamp recordings in hindbrain slices, to demonstrate the role of calcium-induced calcium release in spontaneous miniature outward currents (Yaeger and Coddington 2018). However, to our knowledge no patch-clamp recording of brain region located rostrally to the hindbrain were done in salamanders.

Here, we describe how to generate salamander brain slices, how to visualize the neurons and how to perform patch-clamp recordings. We present the first patch-clamp recordings of neurons in the diencephalon and telencephalon in salamander brain slices. This method should be useful to identify mechanisms at the cellular level in salamander brain microcircuits.

## 2. Material and Methods

### 2.1. Ethics statement

The procedures conformed to the guidelines of the Canadian Council on Animal Care and were approved by the animal care and use committees of the Université de Sherbrooke (QC, Canada). Care was taken to minimize the number of animals used and their suffering.

### 2.2. Animals

A total of 4 Mexican axolotls (*Ambystoma mexicanum*) purchased from the Ambystoma Genetic Stock Center (University of Kentucky, KY, USA) with snout-vent length ranging from 8 to 12 cm were used for the present study. The animals were kept in aquariums at 17-19°C and fed twice per week with fish pellets.

### 2.3. Identification of brain regions

The striatum (Fig. 2A) was identified as part of the ventrolateral pallium based on salamander telencephalon atlases (Northcutt and Kicliter 1980, ten Donkelaar 1998) and our previous anatomical studies showing that injection of a neural tracer in the striatum retrogradely labels dopaminergic neurons in the posterior tuberculum, located at the border of the diencephalon and mesencephalon (Ryczko et al. 2016b). This ascending dopaminergic pathway is considered analogous to the nigrostriatal pathway in mammals (Ryczko et al. 2016b).

**Figure 1.**
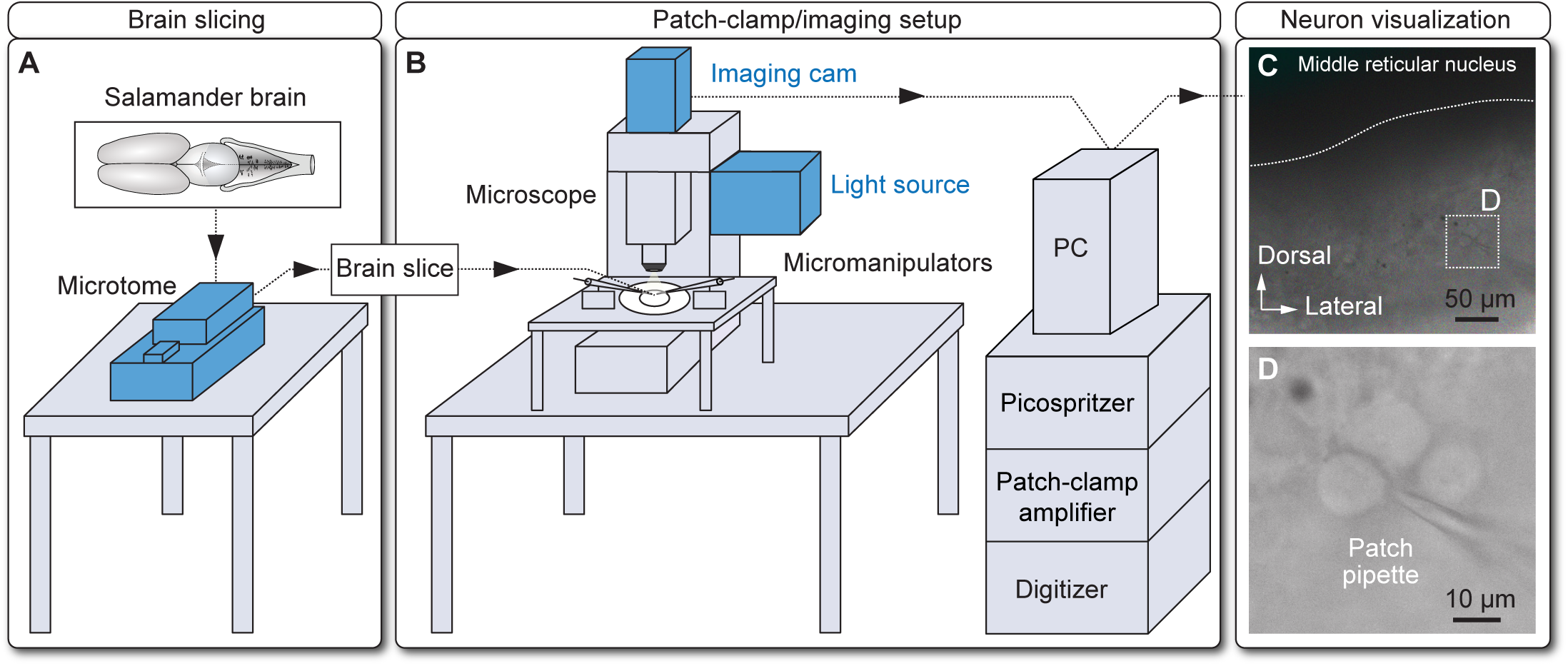
Brain slice preparation and visualization. **A.** Salamander brains were dissected and sliced using a vibrating blade microtome. **B.** Coronal brain slices were placed under the objectives of a microscope coupled with a patch-clamp electrophysiology setup equipped with a Picospritzer pressure microinjector. **C-D.** The imaging camera coupled with DIC components was used to visualize the approach of the patch pipette toward the membrane of neurons for recordings. In C-D, a slice at the level of the middle reticular nucleus is shown. In C, the horizontal white dashed line shows the approximate location of the ventricle border. In D, magnification of the dashed square in C, showing the pipette and a patched neuron.

**Figure 2.**
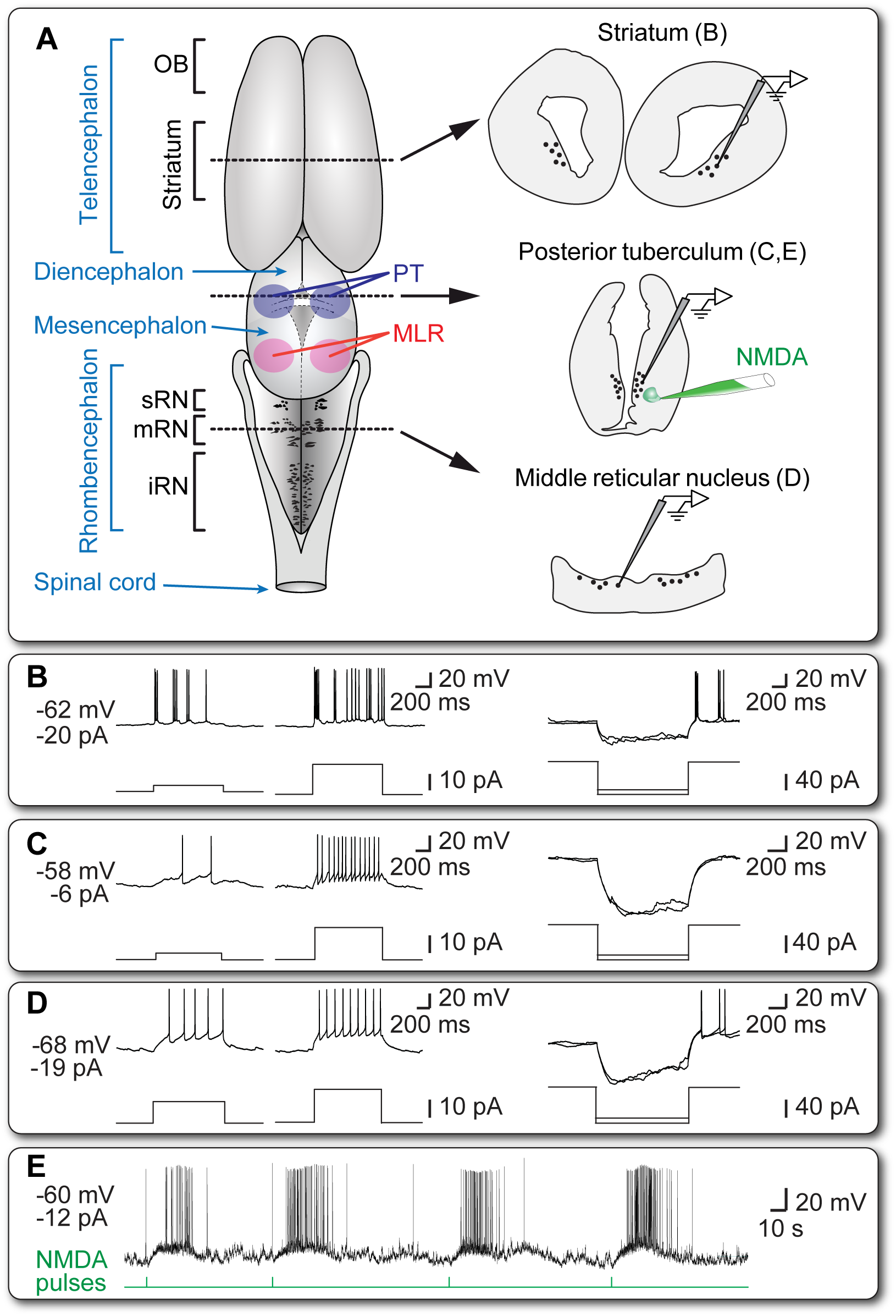
Whole cell patch-clamp recordings in salamander brain slices of telencephalon, diencephalon or rhombencephalon. **A.** Coronal brain slices (350 µm thickness) were obtained from salamander brains at the level of the striatum (telencephalon), posterior tuberculum (PT, diencephalon) and middle reticular nucleus (mRN, rhombencephalon). The black dots illustrate the approximate location of the region targeted for our recordings. **B.** Patch-clamp recordings in current-clamp mode obtained from the striatum (B), posterior tuberculum (C) and middle reticular nucleus (D). Typical neuronal responses to positive current steps (left panels) and negative current steps (right panels) are illustrated. **E.** Four local microinjections of the glutamatergic agonist NMDA (50 ms pulses, 2 mM, 5 psi) applied with a Picospritzer onto the neuron recorded in the posterior tuberculum. Note that each injection evoked a burst of action potentials. In B-E, the membrane potential value and the amount of negative current tonically applied to the cell are indicated on the left part of each panel. Data from B-E were obtained from three different animals. iRN, inferior reticular nucleus; MLR, Mesencephalic Locomotor Region; OB, olfactory bulbs; sRN, superior reticular nucleus.

The posterior tuberculum (Fig. 2A) was identified based on i) our previous anatomical studies showing that dopaminergic neurons in the posterior tuberculum send ascending projections to the striatum (Ryczko et al. 2016b, see also Joven et al. 2018) and descending dopaminergic projections to the Mesencephalic Locomotor Region (Ryczko et al. 2016b); ii) our previous physiological experiments in isolated brains showing that stimulation of the posterior tuberculum evokes large calcium responses in reticulospinal neurons, that relay the locomotor command to the spinal cord (Ryczko et al. 2016b).

The middle reticular nucleus (Fig. 2A) was identified based on: i) previous anatomical studies showing the distribution of reticulospinal neurons in this reticular nucleus (Naujoks-Manteuffel and Manteuffel 1988; Sanchez-Camacho et al., 2001; Chevallier et al. 2004); ii) our previous anatomical results showing that injection of a tracer in this nucleus retrogradely labels neurons in the Mesencephalic Locomotor Region (Ryczko et al. 2016a,b); iii) our previous physiological results showing that reticulospinal neurons in this nucleus respond following stimulation of the Mesencephalic Locomotor Region (Ryczko et al. 2016b); iv) our previous physiological results showing that stimulation of this nucleus generates steering movements in a salamander semi-intact preparation (Ryczko et al. 2016c).

### 2.4. Brain dissection

Animals were anesthetized with tricaine methanesulfonate (MS-222, 200 mg/mL, Sigma) and transferred into a dissection chamber filled with artificial cerebrospinal fluid (aCSF) (in mM: 124 NaCl, 3 KCl, 1.25 KH_2_PO_4_, 1.3 MgSO_4_, 26 NaHCO_3_, 10 Dextrose, and 1.2 CaCl_2_, pH 7.3–7.4, 290–300 mOsmol/kg) bubbled with 95% O_2_ and 5% CO_2_. After evisceration, the skin and muscles were removed carefully to expose the brain and the first two segments of the spinal cord. The meninges were carefully removed, and the cranial nerves were sectioned. A complete transection was done at the level of the first spinal segment, and the brain was removed and dipped in an ice-cold sucrose-based solution (in mM: 3 KCl, 1.25 KH_2_PO4, 4 MgSO_4_, 26 NaHCO_3_, 10 Dextrose, 0.2 CaCl_2_, 219 Sucrose, pH 7.3–7.4, 300-320 mOsmol/kg) bubbled with 95% O_2_ and 5% CO_2_. For slices at the level of the striatum or posterior tuberculum, a transverse section was done with a razor blade at the level of the olfactory bulbs (Fig. 2A). For slices at the level of the brainstem, the transverse section was done at the junction between telencephalon and diencephalon.

### 2.5. Brain slicing

The brain was then glued at the level of the transverse section onto the specimen disk and placed in the slicing chamber filled with the ice-cold sucrose-based solution described above, with the vibratome slicing blade facing the dorsal side of the brain. Coronal slices (350 µm thickness) were prepared with a VT1000S vibrating-blade microtome (also called vibratome, Leica). Blade progression was visually inspected with a stereomicroscope (Leica) installed over the VT1000S. During slicing, high frequency oscillations of the vibratome blade (100 Hz, scale setting “10”) and a slow blade progression (0.15 mm/s, scale setting “3”) were used. To lessen the brain movements evoked by the blade vibrations, a small brush was gently positioned against the ventral side of the brain, at the level of the slice being cut. Slices were then allowed to rest at room temperature for an hour in a chamber filled with aCSF bubbled with 95% O_2_ and 5% CO_2_. Brain slices were carefully placed at the bottom of the recording chamber with the brush under the microscope and secured in place with two platinum wires disposed on the left and right sides of the slice.

### 2.6. Whole-cell patch-clamp recordings

Whole-cell recordings were carried out at room temperature in a recording chamber perfused with aCSF (100 mL/h) bubbled with 95% O_2_ and 5% CO_2_. Neurons were visualized under an Axio Examiner Z1 epifluorescent microscope (Zeiss) equipped with 5× air objective and a 40× water-immersion objective, differential interference contrast (DIC) components, an ORCA-Flash4.0 V3 Digital CMOS camera (Hamamatsu), an halogen light source and a Colibri 7 fluorescent light source (Zeiss). Patch pipettes were pulled from borosilicate glass capillaries (1.0 mm outside diameter, 0.58 mm inside diameter, 1B100F-4, World Precision Instruments) using a P-1000 puller (Sutter Instruments). Pipettes (resistance 6–12 MΩ) were filled with a solution containing (in mM) 140 K-gluconate, 5 NaCl, 2 MgCl_2_, 10 HEPES, 0.5 EGTA, 2 Tris ATP salt, 0.4 Tris GTP salt, pH 7.2–7.3, 280– 300 mOsmol/kg, 0.05 Alexa Fluor 594 or 488, and 0.2% biocytin. Alexa Fluor is useful used to monitor the morphology of the recorded neuron during the experiment. When searching for a cell to record, positive pressure was applied through the glass pipette to prevent it from getting clogged. The neuron membrane was approached with a pipette using a motorized micromanipulator (Sutter instruments). A gigaseal was established by removing the positive pressure. The membrane potential was held at −60 mV, and the membrane patch was suctioned. The pipette resistance and capacitance were compensated electronically, and the neurons were recorded in current-clamp mode. Neurons were discarded when action potential amplitude was less than 40 mV or when the resting membrane potential was too depolarized (>-45 mV). Patch-clamp recordings were performed using a Multiclamp 700B amplifier and a Digidata 1550B digitizer coupled with a computer equipped with PClamp 10 software (Axon Instruments).

### 2.7. Drugs

N-Methyl-D-aspartic acid (NMDA) was purchased from Sigma and diluted to the final concentration of 2 mM in aCSF and applied locally over the recorded neuron with a glass micropipette (tip diameter ∼ 1 μm) using pressure pulses (50 ms duration, 5 psi) applied with a Picospritzer III (Parker).

## 3. Results

### 3.1. Neuron visualization

The brain slices were prepared with the microtome and transferred under the microscope (Fig.1 A-B). The quality of the slice was validated by visual inspection under the microscope with low magnification (5× objective). This step was also used to confirm the rostrocaudal location of the slice based on previous studies and available salamander brain atlases (see section 2.3 in the Methods). Neurons were visualized at higher magnification with the 40× objective coupled with DIC components (Fig. 1C-D). The positive pressure applied through the pipette was essential to navigate through the slice at different depths. As a rule, we chose cells that were deeper than the surface layer, showed a regular cell body shape, without being too dark at their membrane border, and whose membrane clearly displayed a reversible pressure-evoked invagination when the patch pipette was approached.

### 3.2. Whole-cell patch-clamp recordings

We performed whole-cell recordings in current-clamp mode in neurons located in the striatum (Fig. 2A,B) posterior tuberculum (Fig. 2A,C-E) and middle reticular nucleus (Fig. 2A,D). A total of 8 neurons were patched out of the 4 animals used. Stable recordings were obtained in 4 neurons from 3 animals. When applying increments of positive currents through the patch electrode, these neurons responded by spiking action potentials (> 40 mV) in the striatum (Fig. 2B), posterior tuberculum (Fig. 2C) and middle reticular nucleus (Fig. 2D). Increasing the intensity of positive current steps increased spiking frequency (Fig. 2B-D). Applying negative currents evoked membrane potential hyperpolarization in neurons from the three regions (Fig. 2B-D), sometimes followed by a small depolarization during the application of the negative current (“sag”) (e.g. Fig. 2D, right panel). This usually indicates the presence of a hyperpolarization-activated depolarizing current, such as the *I*_*h*_, as documented in salamander motoneurons (Chevallier et al. 2006). Some neurons displayed a post inhibitory rebound and post inhibitory spiking (Fig. 2B,D, right panels). The other 4 patched neurons were discarded because of either unstable membrane potential despite strong negative currents tonically applied to the cell (−80 to −220 pA), rapid cell loss, or small action potential amplitude (< 40 mV).

Next, we determined whether local microinjection of drugs over the recorded neuron could be used to evoke reproductible spiking responses. Repeated microinjections of a glutamatergic agonist (NMDA 2 mM, 50 ms pulse, 5 psi) evoked a consistent burst of action potentials in the neuron recorded in the posterior tuberculum (Fig. 2E), suggesting that these neurons express functional glutamatergic receptors. This is consistent with previous observations showing that glutamate microinjections evoke calcium responses in reticulospinal neurons recorded with calcium imaging in isolated salamander brainstem preparations (Ryczko et al. 2016a).

## 4. Discussion

In the present study we developed the use of patch-clamp recordings of neurons recorded from brain slices of the telencephalon, diencephalon and rhombencephalon of salamanders. We show that microinjections of drugs can be used to design experiments aiming at investigating cellular mechanisms in brain slices.

Previously, no patch-clamp recordings were done in salamander brain regions higher than the spinal cord (Chevallier et al. 2006) and hindbrain (Yaeger and Coddington 2018). To our knowledge, we provide the first whole-cell recordings in the salamander diencephalon and telencephalon. A comprehensive study of the electrophysiological properties of these neurons was not within the scope of the present study. However, such characterization is being carried out in our laboratory and will be reported in the future.

The recent studies have established that the brain and spinal cord of salamanders show striking similarities with that of other vertebrates, including mammals (Ryczko et al. 2016a,b,c). Together with the recent study of Yaeger and Coddington (2018), our present study demonstrates that cellular mechanisms can be studied in any brain area of salamanders using brain slices as classically done in rodents. This approach is an important addition to the diversity of preparations available in salamander neuroscience research, including isolated spinal cords (Ryczko et al. 2015), isolated brains (Ryczko et al. 2016a,b), and semi-intact preparations (Ryczko et al. 2016c). Together with the recent expansion of the genetic toolbox (e.g. Joven et al. 2018), and the use of modeling and robotics (Ijspeert et al. 2007), future work should open new horizons in the understanding of intact and regenerated tetrapod neural circuits using the salamander as an animal model.

## Conflict of interest

The authors declare no competing financial interests.

## Acknowledgments

This work was supported by the Natural Sciences and Engineering Research Council of Canada (RGPIN-2017-05522 and RTI-2019- 00628 to D.R.); the Canadian Institutes of Health Research (407083 to D.R.); the Fonds de la Recherche - Québec (FRQS Junior 1 awards 34920 and 36772 to D.R.); the Centre de Recherche du Centre Hospitalier Universitaire de Sherbrooke (CHUS); the fonds Jean-Luc Mongrain de la fondation du CHUS; the Faculté de médecine et des sciences de la santé; the Centre d’excellence en Neurosciences de l’Université de Sherbrooke.

## Notes

### Competing Interest Statement

The authors have declared no competing interest.

